## Supplemental Files for "Visualizing Molecules of Functional *Human* Profilin"

### Supplemental Information

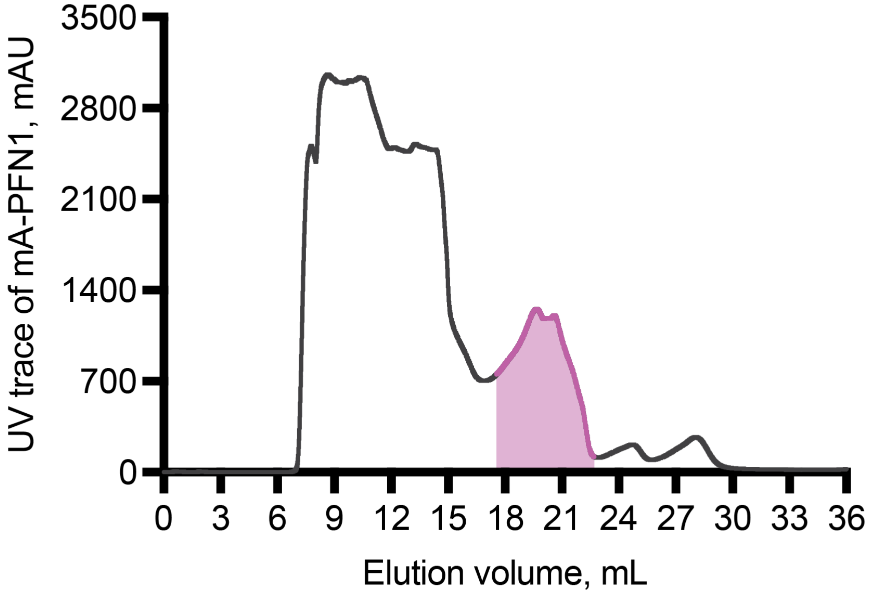


**Figure S1. Gel filtration trace of mApple-profilin-1 (mAp-PFN1).** Trace of mAp-PFN1 (pink) elution from gel filtration column.

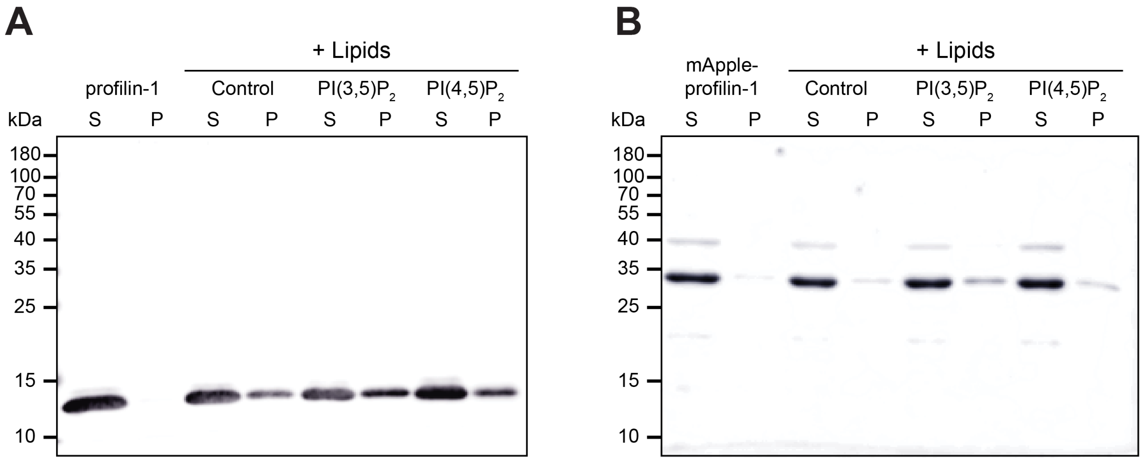


**Figure S2. Full blots associated with tagged PFN1 binding phosphoinositide (PIP)-lipids.** **(A)** Supernatant and pellet samples from liposome pelleting assays in Figure 2B. **(B)** Blot from assay in (A) with mAp-PFN1. Blots were probed with anti-PFN1 antibody (1:5,000; SantaCruz 137235, clone B-10) paired with goat anti-mouse:IRDye 800CW (1:10,000; LI-COR Biosciences 926-32210).

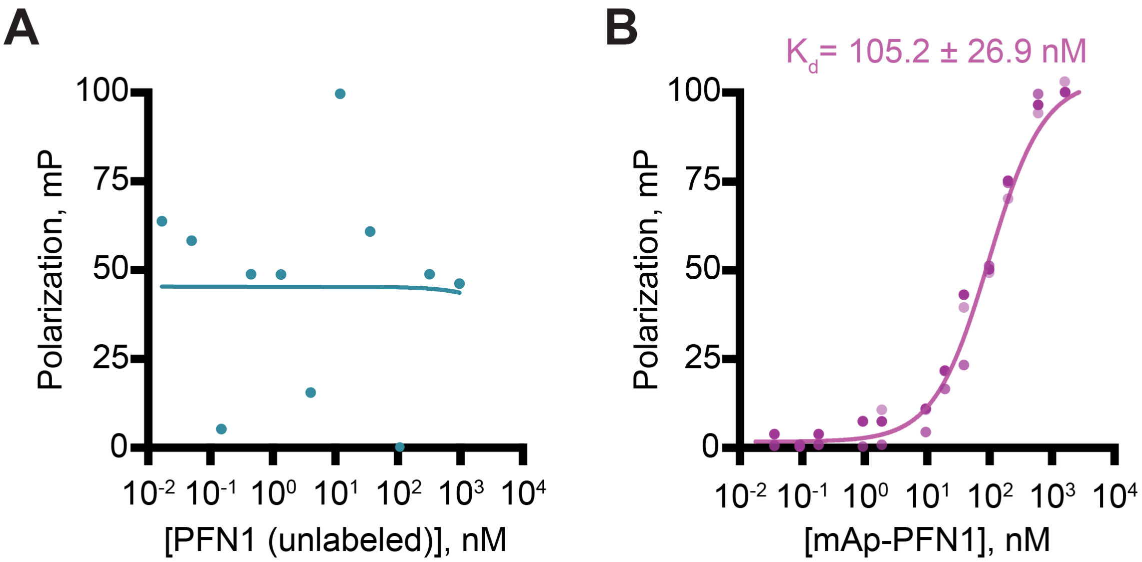


**Figure S3. mAp-PFN1 binds Oregon Green (OG)-actin monomers and is suitable for fluorescence-based binding assays.** **(A)** Fluorescence polarization of 10 nM OG-actin mixed with concentrations of PFN1. PFN1 did not elicit a change in polarization ($n = 1$). **(B)** Polarization of 10 nM actin (unlabeled) with mAp-PFN1 bound actin equivalent to PFN1 (Figure 3B) ($n = 3$).

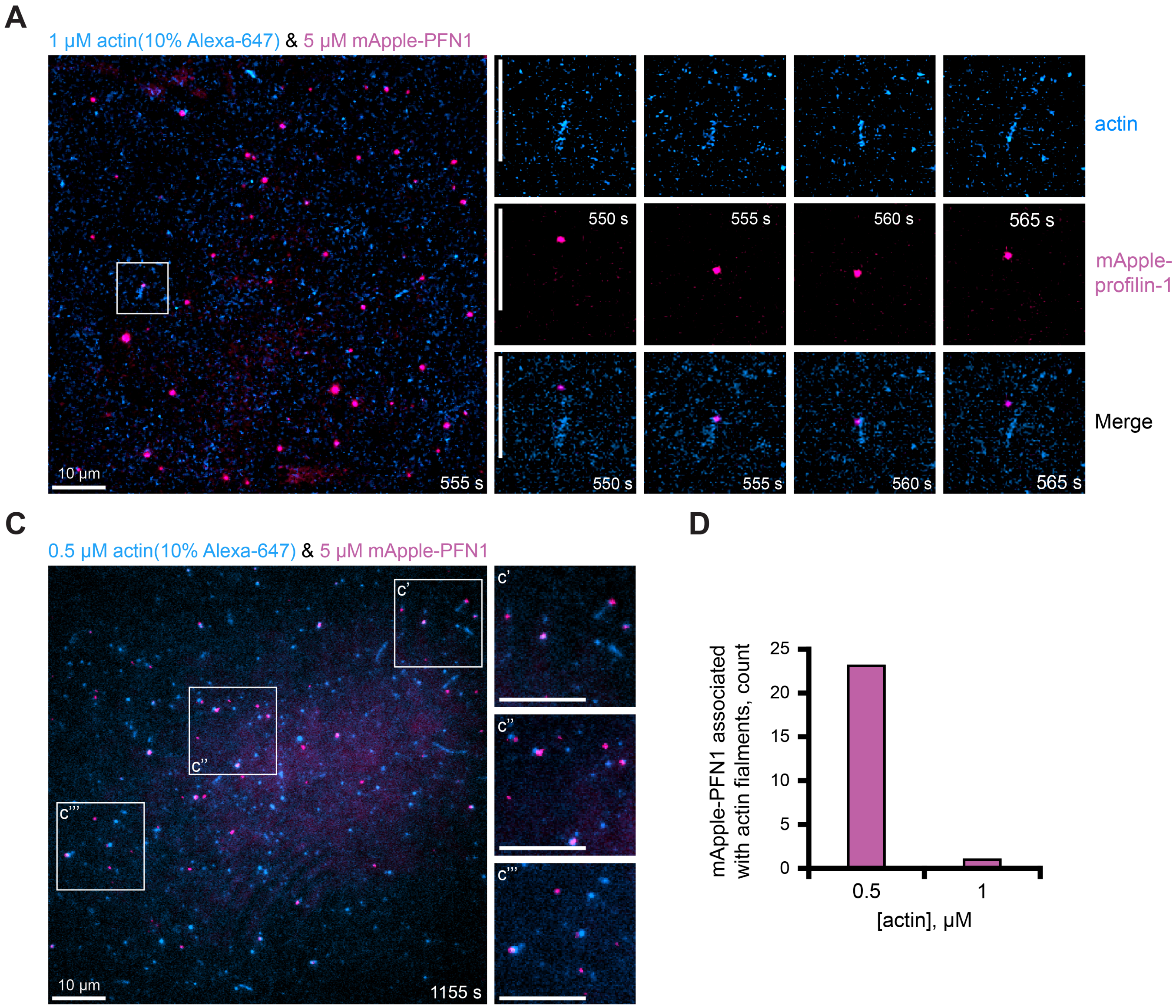


**Figure S4. Localization of mAp-PFN1 with actin filaments in vitro.** **(A)** Image and montage from a TIRF assay containing: 1 μM actin (10% Alexa-647-labeled; 0.6 nM biotin-actin) (blue), and 5 μM mAp-PFN1 (pink). **(B)** Magnified view of (A). **(C)** Image of 0.5 μM actin (10% Alexa-647-labeled; 0.6 nM biotin-actin) (blue) and 5 μM mAp-PFN1. **(D)** Quantification of mAp-PFN1 with actin. Scale bars, 10 μm.

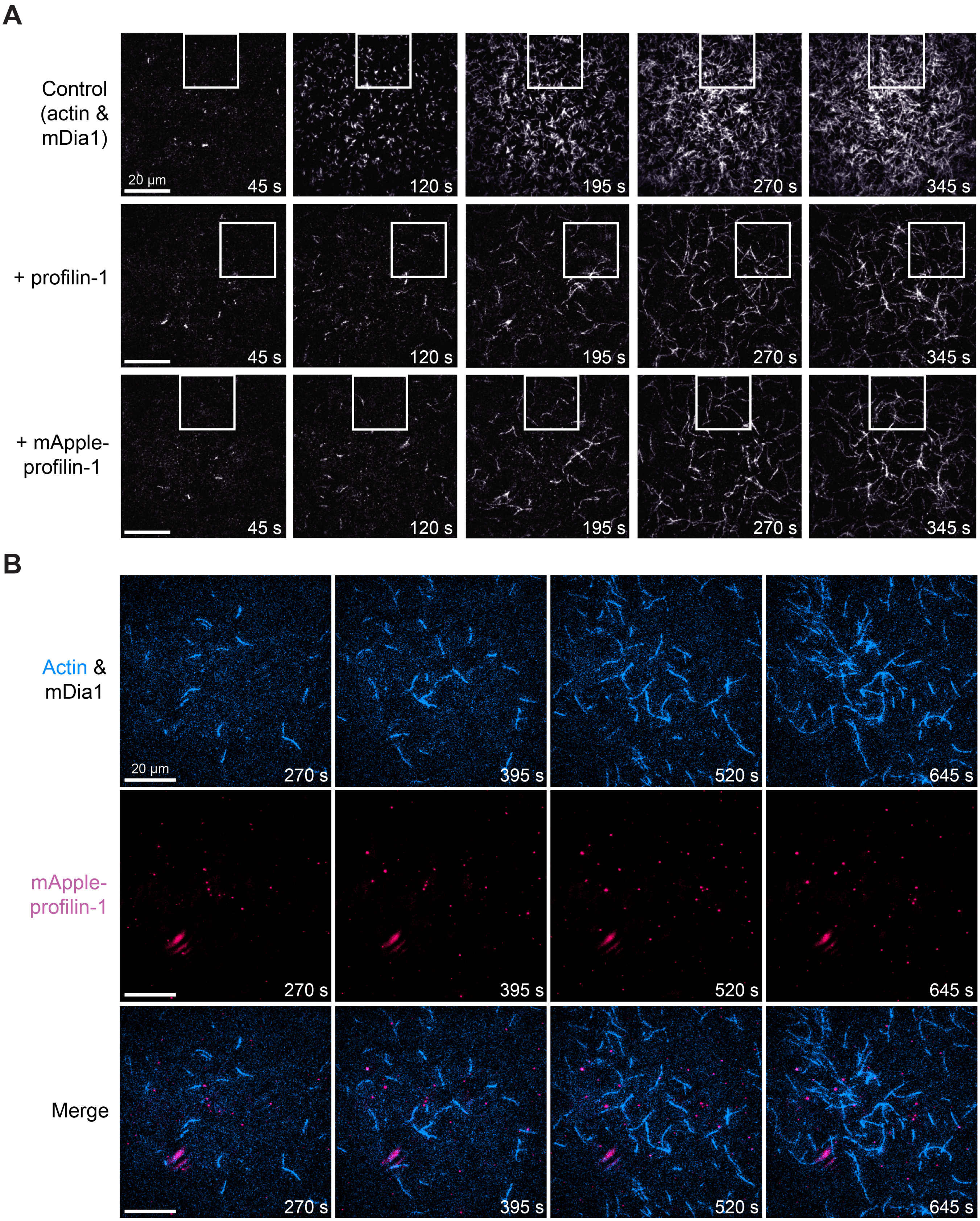


**Figure S5. Full views of mAp-PFN1 on formin-mediated actin assembly. (A)** Full views formin-mediated actin polymerization montage in Figure 4B (white boxes). **(B)** Multi-color TIRF montage containing: 1 µM actin (10% Alexa-647-labeled; 0.6 nM biotin-actin), 25 nM mDia1(FH1-C) and 5 µM mAp-PFN1. Scale bars, 20 µm.

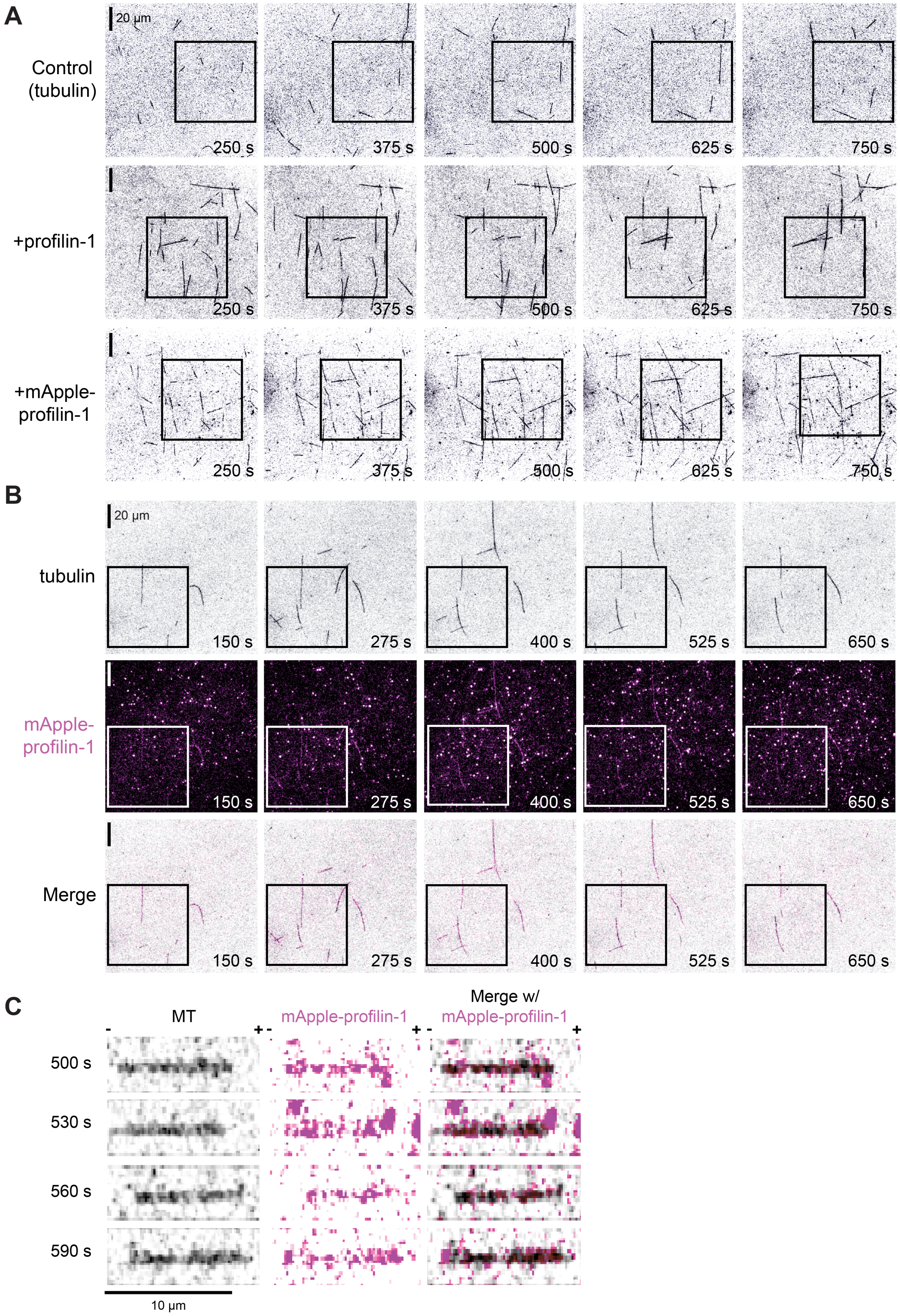


**Figure S6. Additional views of the effects of mAp-PFN1 on microtubule dynamics. (A)** Full views of montages present in Figure 5A (black boxes). **(B)** Full view of multi-color time lapse TIRF montage present in Figure 5H (black boxes). Scale bars, 20 µm. **(C)** Montage of a microtubule (black), mAp-PFN1 (pink) and merge from Figure 5I. Scale bar, 10 µm. + and - indicate microtubule polarity.

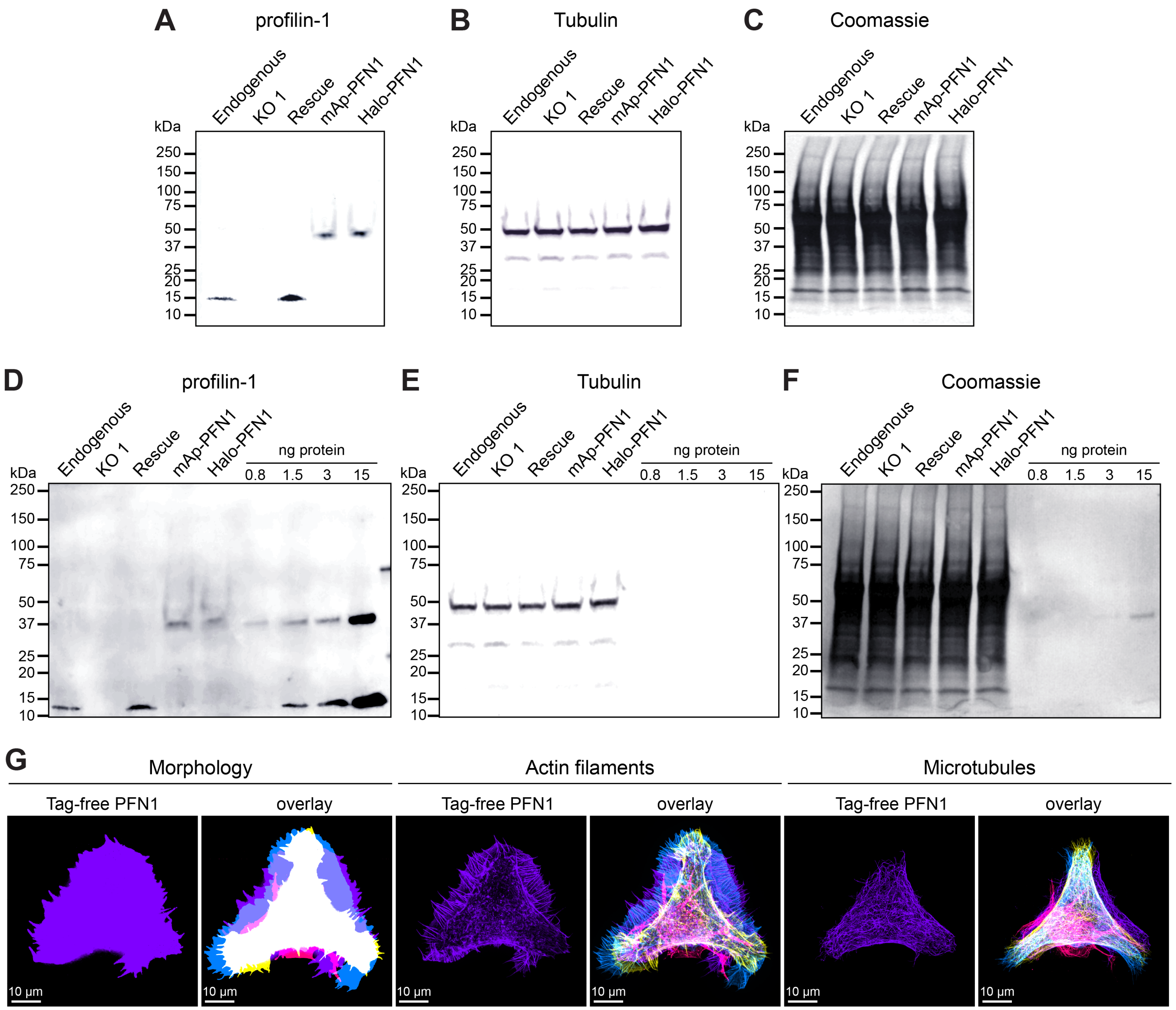


**Figure S7. Full blots used to determine PFN1 levels in Neuroblastoma-2a (N2a) cells.** Blots confirming knockout or rescue of PFN1 as in Figure 6A. **(A)** Blots were probed with anti-PFN1 antibody (1:3,500; SantaCruz 137235, clone B-10) paired with goat anti-mouse:IRDye 800CW secondary antibody (1:5,000; LI-COR Biosciences 926-32210) and **(B)** anti-α-tubulin primary (1:10,000; Abcam 18251) paired with donkey anti-rabbit: IRDye 680RD (1:20,000; LI-COR Biosciences 926-68073). **(C)** Coomassie stained membrane from (A). (D) Example blot used to determine PFN1 concentration N2a cells. Blot probed as in (A-C) for (D) PFN1 and **(E)** α-tubulin, and **(F)** Coomassie stained. **(G)** Morphology measurements related to Figure 6 for tag-free PFN1.

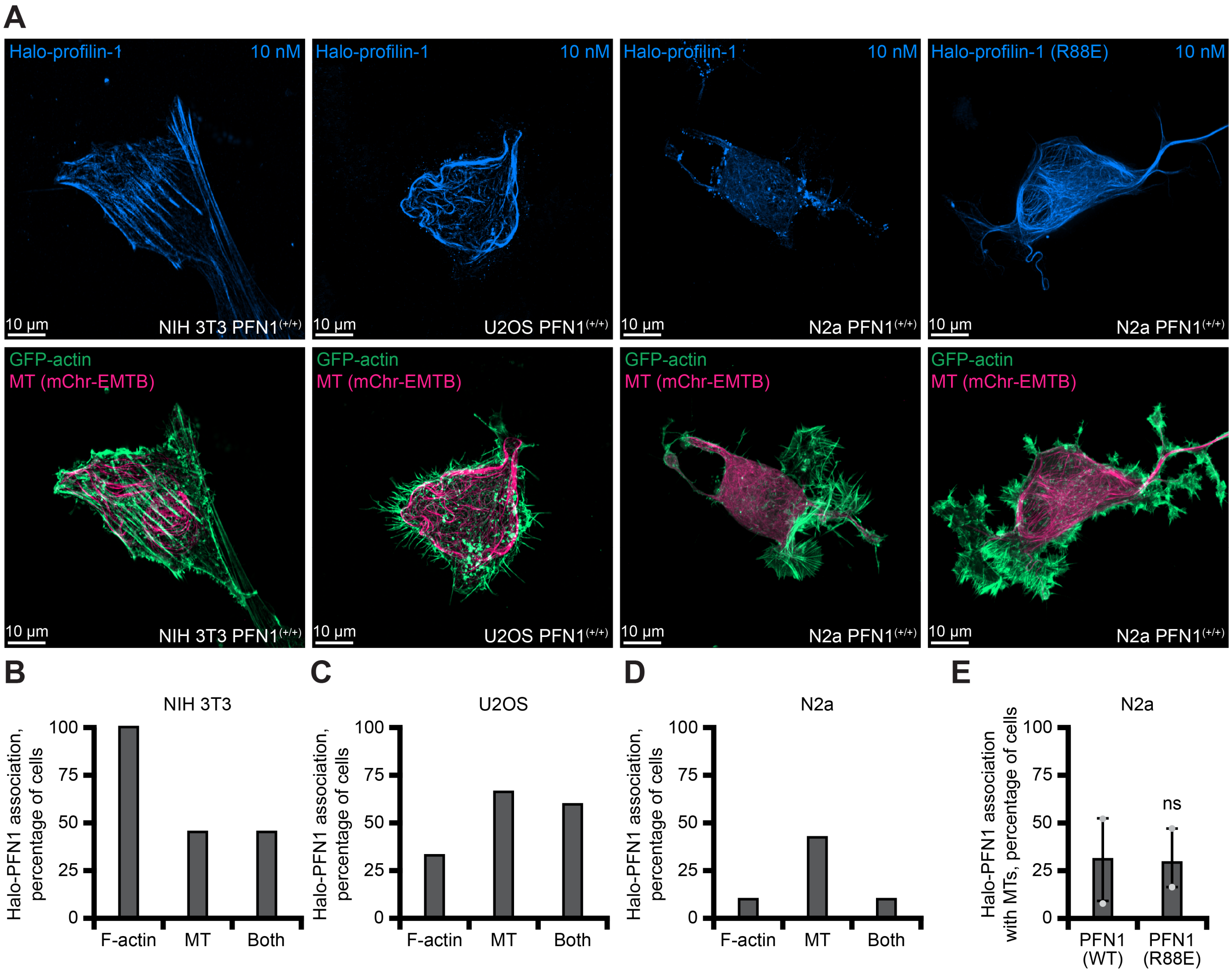


**Figure S8. Localization of Halo-PFN1 in different cell types.** **(A)** Maximum intensity images of different PFN1(+/+) cells transiently expressing GFP-actin (green), a marker for microtubules (EMTB-2×mCherry) (pink), and either Halo-PFN1 (Halo-PFN1) or Halo-PFN1(R88E) (light blue), visualized with 10 nM JF-646 ligand. Scale bars, 10 µm. **(B-D)** Quantification of Halo-PFN1-microtubule overlap from different cell types from (A). **(E)** No difference was found Halo-PFN1 or Halo-PFN1(R88E)-microtubule colocalization.

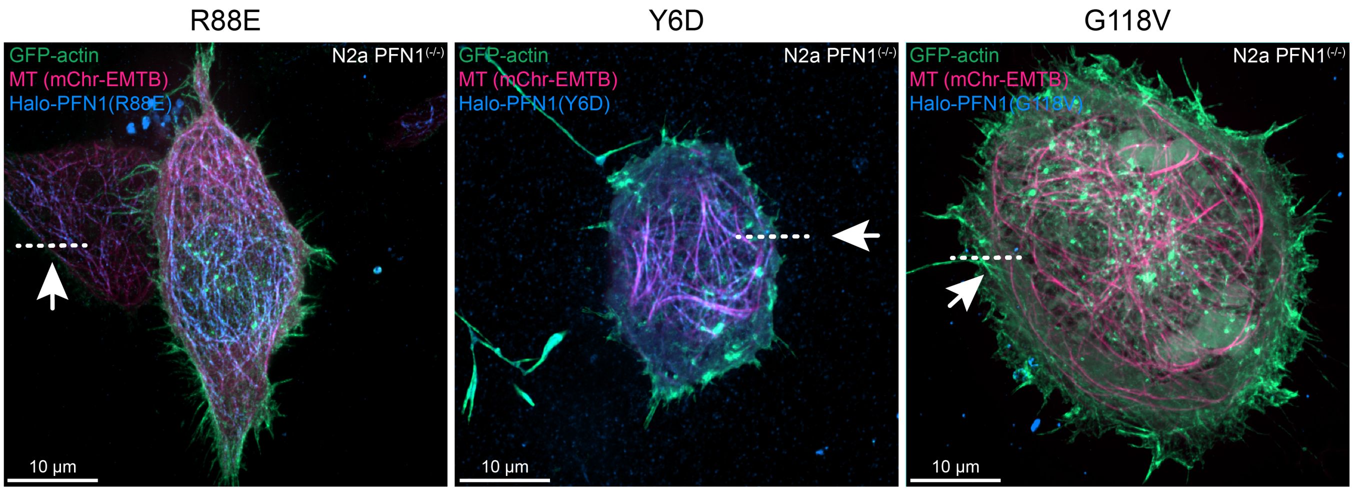


**Figure S9. Live cell localization of constructs of Halo-PFN1.** View of kymograph line drawn on cells expressing Halo-PFN1 mutants in Figure 8D. Scale bars, 10 μm.

### Supplemental movies

**Supplemental Movie 1. TIRF movie of PFN1 or mAp-PFN1 on actin assembly.** Reaction contains: 1 µM actin (20% OG-labeled; 0.6 nM biotin-actin) with buffer (control) or 3 µM PFN1 or mAp-PFN1. Playback,10 fps. Scale bars, 10 µm.

**Supplemental Movie 2. TIRF movie of mAp-PFN1 on actin assembly.** Reaction contains: 1 µM actin (20% OG-labeled; 0.6 nM biotin-actin) (cyan) and 3 µM mAp-PFN1 (pink). Box corresponds to Figure S4A inset. Playback, 10 fps. Scale bars, 10 µm.

**Supplemental Movie 3. TIRF movie of PFN1 or mAp-PFN1 on formin-mediated actin assembly.** Reaction contains: 1 µM actin monomers (10% Alexa-647-labeled; 0.6 nM biotin-actin) and 5 µM PFN1 or mAp-PFN1. Box corresponds to montage in Figures 4B and S5A. Playback, 10 fps. Scale bars, 10 µm.

**Supplemental Movie 4. TIRF movie of mAp-PFN1 on formin-mediated actin assembly.** Reaction contains: 1 µM actin (10% Alexa-647-labeled; 0.6 nM biotin-actin) (cyan) and 5 µM mAp-PFN1 (pink). Playback,10 fps. Scale bars, 10 µm.

**Supplemental Movie 5. TIRF movie of PFN1 or mAp-PFN1 on microtubules.** Reaction contains: 647-biotinylated-GMP-CCP microtubule seeds (not shown), 10 µM tubulin (5% HiLyte-488) in buffer or 5 µM profilin or mAp-PFN1. Box corresponds to Figure 5A and S6A. Playback, 10 fps. Scale bars, 20 µm.

**Supplemental Movie 6. TIRF movie of mAp-PFN1 on microtubules.** Reaction contains: 647-biotinylated-GMP-CCP microtubule seeds (not shown), 10 µM free tubulin (5% HiLyte-488) (black) and 5 µM mAp-PFN1 (pink). Box corresponds to montage inset from Figure 5H and S6B. Playback 10 fps. Scale bars, 20 µm.

**Supplemental Movie 7. mAp-PFN1 transiently associates with the microtubule lattice.** Reaction contains: 647-biotinylated-GMP-CCP microtubule seeds (not shown), 10 µM free tubulin (5% HiLyte-488) (black) and 5 µM mAp-PFN1 (pink). + and -, microtubule polarity. Playback, 10 fps. Scale bar, 10 µm.

**Supplemental Movie 8. Halo-PFN1 dynamics in live N2a cells.** 4D-spinning disk confocal movie of cells transiently expressing markers for actin (GFP-actin; cyan), microtubules (EMTB; yellow), and Halo-PFN1 plasmids labeled with 10 nM JF-646 (magenta). Playback, 10 fps. Scale bar, 10 µm.
